# Automatic Discovery of Optimal Discrete Character Models

**DOI:** 10.1101/2024.11.15.623760

**Authors:** James D. Boyko

## Abstract

Modeling discrete character evolution in a Markovian framework has become common practice in phylogenetic comparative methods. The increasing size and complexity of these models reflects a trend of analyses to include more taxa and more discrete characters. However, as complexity of the models increase, so do the number of potential model structures and number of estimable parameters, making it nearly impossible to consider all modeling options for a given dataset. To overcome this issue, I apply a combination of regularization and simulated annealing to models of discrete character evolution. This allows for the automatic searching and optimization across different model structures without user specification. I test this framework under several simulation scenarios including hidden rates and multiple discrete characters. The results indicate that regularized models significantly outperform traditional approaches, yielding a far lower variance and a nearly tenfold reduction in the overall error of parameter estimates in the most extreme scenarios. I illustrate the power of automatic model selection by revisiting the ancestral state estimation of concealed ovulation and mating systems in Old World monkeys. Using the dredge algorithm, I discover a previously unexamined model structure which has both better statistical performance and a differing ancestral state reconstruction when compared to default model sets. In general, these results highlight the dangers of an over-reliance on default model sets. The combination of automatic model selection and regularization help overcome problems of over-parameterization, and these results demonstrate that when inferences are drawn from a larger model space, they can be both more statistically robust and biologically realistic.

---

Multistate discrete character models are now widely applied in phylogenetic comparative methods (PCMs). These models, which were initially limited to relatively few characters and simple processes, have been expanded in an attempt to incorporate various biological processes and sources of variation. Complexity has been introduced to these models through correlated character evolution (Pagel 1994), hidden rate variation (Beaulieu et al. 2013; Boyko and Beaulieu 2021), state-dependent speciation and extinction (SSE) (Maddison et al. 2007; Beaulieu and O’Meara 2016), and the incorporation of continuous character information (May and Moore 2020; Boyko et al. 2023).

This increasing complexity has also led to higher generality. For example, hidden Markov models (HMMs) which were initially introduced to model two binary characters (Beaulieu et al. 2013), have been expanded to allow for any number of characters, observed states, and hidden states (Boyko and Beaulieu 2021). However, while this additional flexibility allows for customization of the phylogenetic comparative method to the system at hand, the large model space can make it challenging for biologists to find the most appropriate model for their particular dataset. The typical approach, multi-model inference framework (Burnham and Anderson 2002), has a biologist decide on a set of potentially realistic models. The relative support for each model is then evaluated based on their fit to the data using information criteria such as Akaike Information Criterion (AIC). This approach is powerful because it allows for model averaging, where inferences are made based on a weighted average of the models included in the set (Burnham and Anderson 2002). However, the effectiveness of multi-model inference relies heavily on the appropriateness of the model set. Several PCMs have been criticized for high false positive rates due to the exclusion of the “correct” null hypothesis (Rabosky and Goldberg 2015). Although these criticisms have been addressed by introducing new model structures to serve as better null hypotheses (Beaulieu and O’Meara 2016; Boyko and Beaulieu 2023), the problem, when recast as a failure to include a complete model set (Boyko and Beaulieu 2023), suggests that there is still a vast space of unexplored model structures.

Often the model set chosen for discrete character evolution are nested, with the difference being which parameters are variable, and which are fixed. For example, for a two-character binary state dataset, the difference between a correlated model (k=8) and the independent model (k=4) is whether changes in the focal character depend on the state of the background character (Pagel and Meade 2006). In the independent model parameters representing this dependency process are fixed to be equal, while in the correlated model they are freely estimated (Pagel and Meade 2006; Boyko and Beaulieu 2023). In this way, one can think of discrete models adding complexity as adding parameters to increase “biological realism” and represent new processes or relationships between variables.

This increasing model complexity then leads to another, more technical, challenge. As the number of estimable parameters grows it reaches a point where the number of parameters (k) approaches the number of taxa (N). However, the rate at which the number of parameters increase often surpasses the rate at which information can be gained through data (Felsenstein 2012). In the case of correlated discrete character models, this is because each additional character requires considering its relationship to all other traits. For instance, the most complex discrete model for a single binary character has 2 parameters, while the most complex model with two binary characters has 12 parameters (k=8 if excluding dual transitions), and the most complex model with three binary characters has 56 parameters (k=26 if excluding dual transitions; Fig. 1). In each instance we have added a single character, but because we must consider the new character’s relationship to all other existing characters, the model complexity (as measured by the number of parameters) outpaces the potential information gained from the new data. This is problematic because although Maximum likelihood methods are consistent estimators when N >> k, their performance deteriorates as models become more parameter-rich, leading to unreliable and biased parameter estimates (Huelsenbeck et al. 2001). However, for complex models with a finite data, it is unlikely that all parameters will be essential or necessary to best model the data (Lemey et al. 2009; Gelman et al. 2013).

**Figure 1.**
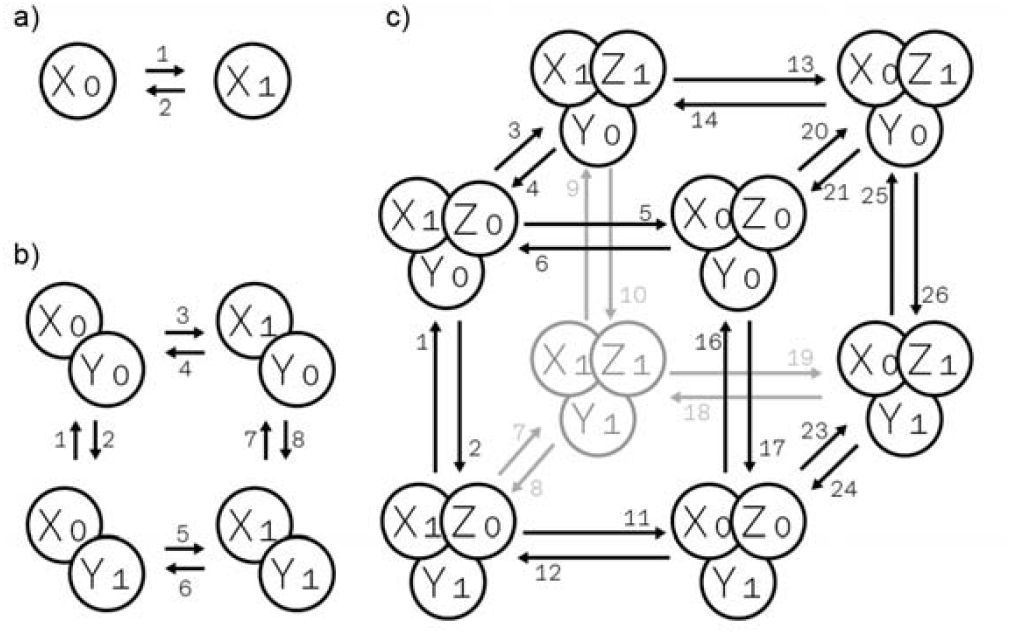
Visualization of the exponential increase in model complexity as additional binary characters are introduced. (a) A single binary character with two states (k=2). (b) Two binary characters, each with two states (k=8, excluding dual transitions). (c) Three binary characters, each with two states (k=26, excluding dual transitions). Adding characters exponentially increases the number of parameters due to the need to account for relationships between all characters. The disallowance of simultaneous multistate transitions keep the models from being even more complex.

Here, I address this challenge by introducing an approach for automatic model exploration using regularization and simulated annealing. Regularization techniques, such as L1 (lasso) and L2 (ridge), constrain the magnitude of parameter estimates and encourage simpler, more generalizable models (Hoerl and Kennard 1970; Tibshirani 1996). By incorporating regularization, we can balance model complexity and goodness-of-fit, reducing the risk of overfitting and improving the stability of parameter estimates. Additionally, to search the set of possible models, I implement a simulated annealing routine that stochastically proposes and accepts model modifications according to a temperature-controlled optimization schedule. This approach is most similar to Pagel and Meade (2006), who introduced a reversible-jump Markov chain Monte Carlo (RJMCMC) to search across models with different numbers of parameters. Their method explores the model space by proposing merge/split and augment/reduce “moves”, which effectively add or remove parameters during sampling. Models are visited in proportion to their posterior probability and produce a distribution of supported models rather than selecting a single best model. While RJMCMC provides a rigorous approach to model selection, it is computationally intensive and currently limited in scope as existing implementations only support two binary characters and do not accommodate hidden states. My framework addresses these limitations by integrating simulated annealing and regularization within corHMM (Beaulieu et al. 2022) implemented as the function corHMMDredge.

To test the dredge framework, I conduct an extensive set of simulations to explore the biasvariance trade-off associated with regularization. Under regularization, it is expected that models will have increased generality and lower variance. I test these expectations by examining regularized model’s accuracy of parameter estimates. Additionally, I perform a more detailed simulation test on a subset of historically important discrete models, such as the hidden rates model (Beaulieu et al. 2013) and correlated character evolution model (Pagel 1999; Pagel and Meade 2006; Boyko and Beaulieu 2021). Finally, as a case study, I reanalyze the dataset from Pagel and Meade (2006) testing whether female Old World monkey estrus advertisement is associated with multimale mating systems. I use this case study to demonstrate the dredge framework, guiding users through each step of the process including cross-validation and uncertainty estimation.

## Methods

### Regularization and Simulated Annealing

The likelihood of a discrete character model with its underlying framework as a continuous time Markov chain (CTMC) is calculated as *L*= *P*(*D*|*Q, Φ, π*), which is the probability of observing the data (D) given an instantaneous rate matrix (Q), root state frequency vector (π), and a phylogeny with a fixed topology and set of branch lengths (Φ). The data consist of the observed character states (S) at the tips of the phylogeny, while the rate matrix contains the rates of character state transitions (q_ij_). The likelihood function is then computed by integrating the product of transition probabilities along the branches of the phylogeny (Felsenstein 1981, 2004; Pagel 1994; Lewis 2001). Though not mathematically necessary, it may be useful to consider a mapping matrix (M; also referred to as an index matrix (Revel 2024)) which gives the structure of the discrete model by indicating which transition rates are estimated and/or fixed to be equal (FitzJohn, 2012; Beaulieu & O’Meara, 2016). For example, if we consider a simple binary character with states 1 and 2, the instantaneous rate matrix is given by 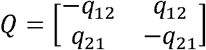. However, the mapping matrix can specify several alternative model structures. If 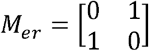, then *q*_12_ = *q*_21_ and we have what is commonly known as the “equal rates” model. An “all rates different” model can be indicated by 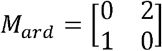, where *q*_12_ ≠ *q*_21_and a “unidirectional” model can be constructed via 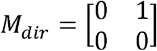, where *q*_21_ = 0. In this specific example, we have models of 1 (“equal rates” and “unidirectional”) or 2 (“all rates different”) estimable parameters. To determine which model structure is optimal (best balances goodness-of-fit and complexity) for the dataset, we can compute the maximum likelihood estimate (MLE) for each model structure and compare them using a information criterion such as AIC (Akaike 1974; Burnham and Anderson 2002).

This framework has been successfully applied for many years within PCMs (O’Meara 2012), but growing model complexity (e.g. Fig. 1) is making it untenable for users to define all possible relevant model structures or for method developers to construct a complete set of default models. As such, regularization techniques may be a necessary step in discovering optimal model structures for large and complex datasets. In corHMMDredge, I incorporate three regularization approaches, two of which are analogous to *l1* and *l2* regularization (Hastie et al. 2015), and the third is based on the null expectation that transition rates are all equal (*er*; (Zhou et al. 2024). Specifically, the regularized likelihoods are defined as:

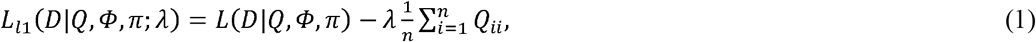

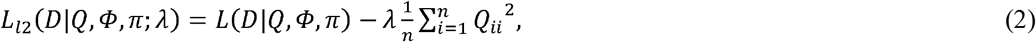

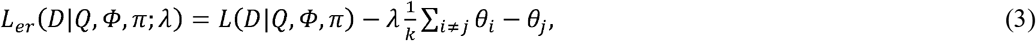

where *L*(*D*|*Q*, Φ, π)denotes the standard likelihood, 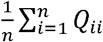 is the elementwise mean of the diagonal of the instantaneous rate matrix, 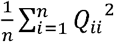 is the squared elementwise mean of the diagonal of the instantaneous rate matrix, 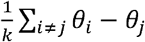 is the mean pairwise distances between all freely estimated parameters in their raw units, *k* is the number of freely estimate parameters, and *λ* is a hyper-parameter that adjusts the severity of the penalty, ranging from 0 (no regularization) to 1 (full penalization). This penalization scheme uses the mean rather than the sum of the parameter values because the number of transition rates is a function of the number of possible discrete states and using the sum would cause the penalization term to be a function of the number of states rather than the overall complexity of the model (the number of parameters). It is important to note that *λ* cannot be jointly estimated alongside the parameters via the maximized likelihood (Clavel et al. 2019). Instead, a form of cross-validation is necessary to tune *λ* (see *Phylogenetically informed k-fold cross validation*). This scheme is similar to Sanderson’s penalized likelihood approach for estimating chronograms from phylogenetic trees (Sanderson 2002). Under penalized likelihood, the penalty is based on differences of rates and is essentially a regularization approach. Furthermore, the Sanderson method includes cross validation to estimate the equivalent of the *λ* hyper-parameter (which he also labeled *λ*, the smoothing parameter).

The second component of the dredge algorithm involves a method to explore and assess alternative model structures (Fig. 2). For this, I implemented a simulated annealing heuristic search algorithm. Simulated annealing is a probabilistic optimization technique inspired by the process of annealing in metallurgy, where a material is heated and then slowly cooled. The algorithm aims to find a model that minimizes a given information criterion (e.g., AIC, BIC) and given enough time is guaranteed to find the optimal solution for a discrete set (Kirkpatrick et al. 1983). This approach is similar to Pagel and Meade (2006), who also use a Metropolis-Hastings sampling scheme to explore alternative models in a Bayesian framework. While their method samples from the posterior distribution, corHMMDredge optimizes each proposed model structure using a penalized likelihood. In both approaches, poor-fitting models can still be temporarily accepted to escape local optima. The search is performed independently for each number of hidden rate categories, from the initial category up to max.rate.cat. Models are evaluated through an iterative process governed by a “temperature” parameter, T, which starts high and is gradually lowered.

**Figure 2.**
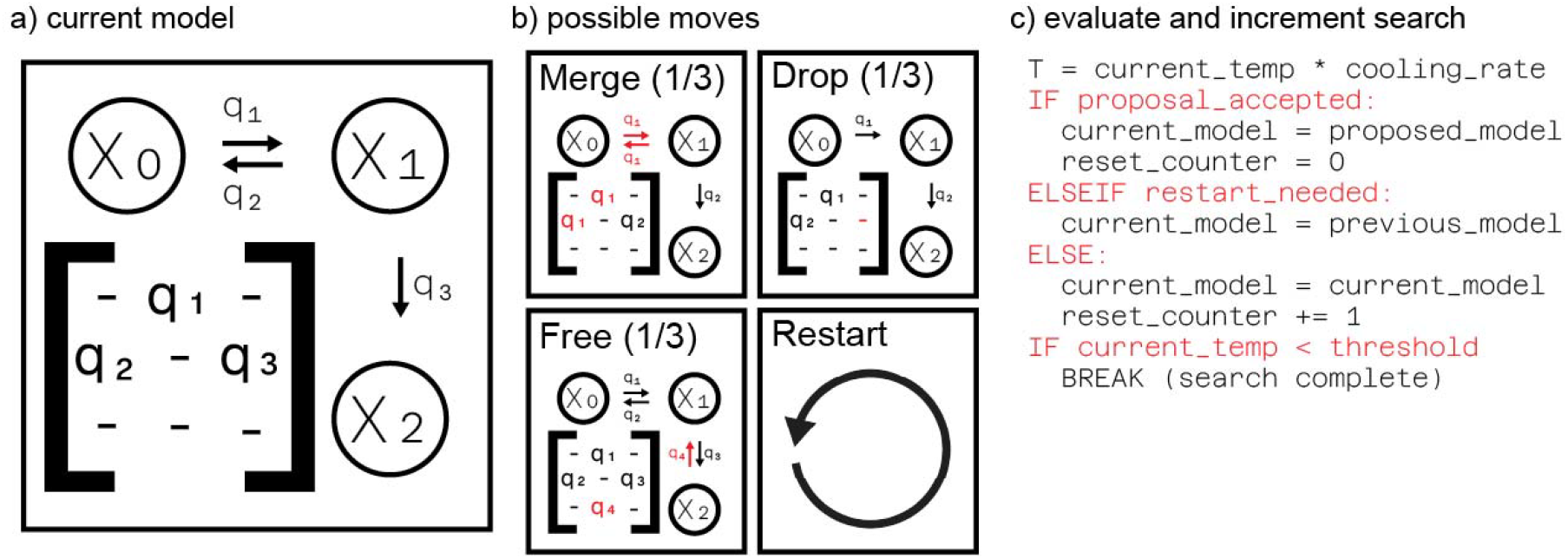
An overview of the simulated annealing algorithm. From a (a) current model, the algorithm proposes a new model by stochastically selecting a move. (b) Possible moves include simplifying the model by merging or dropping parameters or increasing complexity by freeing a constrained parameter. A restart mechanism is also available to escape local optima. (c) The algorithm then decides whether to accept the new model. Better models are always accepted, while worse models may be accepted with a probability that decreases as the search “temperature” cools, allowing the search to converge on an optimal solution.

The search begins with an initial model structure (default is the model with the highest number of free parameters) and its corresponding score. At each iteration, the algorithm proposes a move to a new, neighboring model structure by making a small perturbation to the current mapping matrix. The proposed model is then fit to the data, and its score is calculated. The decision to accept this new model is probabilistic and depends on the change in the information criterion score (ΔE = E_new - E_current) and the current temperature, T. As in Pagel and Meade (2006), if ΔE ≤ 0, the proposed model is an improvement (or equally good), and it is always accepted as the new current model. If ΔE > 0, the proposed model is worse than the current one, but following the Metropolis algorithm (Metropolis et al. 1953), it may still be accepted with a probability P = exp(-ΔE /T). When the temperature T is high, the acceptance probability is also high, allowing the search to freely explore the entire model space and escape local minima. As the algorithm proceeds, T is gradually reduced according to an exponential cooling schedule (T_i+1 = T_i * α, where α is a cooling.rate typically close to 1.0 – default of 0.95). As T approaches zero, the probability of accepting a worse model diminishes, causing the algorithm to behave like a greedy hill-climbing search (Kirkpatrick et al. 1983; Russell and Norvig 2021).

To generate new candidate models, one of three stochastic move types is randomly selected at each iteration: drop, merge, or free. A parameter drop simplifies the model. The algorithm identifies all estimated rate parameters and stochastically selects one to remove, with the probability of selection being inversely proportional to the parameter’s estimated rate (i.e., smaller rates are more likely to be dropped). Second, a parameter merge also simplifies the model by introducing a constraint. The algorithm identifies pairs of parameters with similar estimated rates. A pair is randomly selected, with the probability of selection being inversely proportional to the absolute difference between their rates (i.e., more similar rates are more likely to be merged). The parameters are then constrained to be equal. For hidden Markov models with multiple rate categories, this merging is restricted to occur only among parameters within the same underlying rate class. Finally, a parameter free move increases model complexity. It identifies transitions that are currently constrained (i.e., previously dropped or merged) and randomly selects one to “free” by assigning it a new, unique parameter index. Note that these three move types (drop, merge, free) are conceptually similar to the proposals used in reversible-jump MCMC algorithms (Pagel and Meade 2006). Like Pagel and Meade (2006), corHMMDredge proposes model simplifications or expansions through stochastic rules. However, unlike an RJ-MCMC, corHMMdredge does not rely on priors and marginal likelihoods to navigate model space, instead the dredge framework utilizes heuristics based on rate magnitudes to propose moves and evaluates model quality using information criteria. This should make the method more scalable to high-dimensional model spaces.

Finally, to prevent the search from becoming permanently trapped in a local minimum, a restart mechanism is included. If the search fails to find a new best-fitting model or has not accepted a new model for a predefined number of iterations (default of 20 attempts), the algorithm reverts to a randomly selected, previously accepted (but not necessarily optimal) model from its search history. The simulated annealing process for a given rate category terminates after a maximum number of iterations or when the temperature becomes negligibly small. The algorithm maintains a record of all unique models accepted during the search. Upon completion, any redundant models (i.e., models with identical log-likelihoods and parameter counts but different rate structures) are pruned, and the remaining unique models are ranked by their information criterion scores to provide a set of models for further consideration.

### Phylogenetically informed k-fold cross validation

To optimize *λ* I choose to employ a phylogenetically informed k-fold cross-validation procedure across the tips of the phylogeny. Cross-validation approaches, such as leave-one-out cross-validation (Lartillot 2023), have been shown to be more adequate than alternatives for model selection. Here I implement a k-fold method because it provides a computationally efficient and intuitive framework that directly addresses phylogenetic non-independence through weighted sampling. K-fold cross-validation is a common way to optimize *λ* when using regularization (Hastie et al. 2009). It involves dividing the dataset into k separate subsets (or folds). The model is trained on k-1 of these folds with the remaining fold used as the test set to evaluate the models’ performance. This process is repeated k times, with each of the k folds used exactly once as the test set. The results of these k tests are then averaged to produce a single estimate. In the context of a discrete character model the metric chosen for evaluation is reconstruction of tip state values. Each species within a k-fold is coded as having an unknown state and a reconstruction based on the fitted model is then used for evaluation. The score for a fold is determined by the average Jensen-Shannon divergence between the predicted and actual tip states. Specifically, 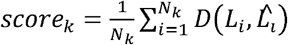 where *N*_*k*_ is the number of tips in the k-th fold, *L*_*i*_ are the observed state likelihoods at the tips of the k-th fold (1 for the observed state and 0 for an unobserved state), 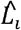 are the predicted state likelihoods for the tips in the k-th fold, and *D* is the Jensen-Shannon (JS) divergence (or any form of divergence more generally). The k scores are then averaged over all folds to obtain a single score for the model. The JS divergence (available in corHMM as the function js_divergence) is chosen because it is a symmetric and bounded measure of difference between predicted and observed probability distributions while being robust to zero probabilities. An advantage of using tip values to measure the model fit is that the phylogenetic structure of the data is unchanged throughout the entire cross-validation procedure because no tips need to be explicitly dropped when fitting the model; they only need to be set to an unknown value. The samples within a given k-fold are chosen without replacement and with probabilities equal to 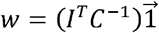 (Rohlf 2001) to ensure phylogenetically even sampling between the k-folds, where w is roughly proportional to the phylogenetic uniqueness of each species.

### Simulation Study I: Bias and variance of parameter estimates

The expected behavior of a regularized model is that the introduction of a penalty term to the likelihood will reduce sensitivity to the specific dataset, resulting in lower variance and greater generalizability at the cost of a systematic error (bias). Though the behavior of regularization techniques is well documented in generalized least squares regression (Hastie et al. 2009), it is less well studied in the context of phylogenetic comparative methods (but see Khabbazian et al. 2016; Clavel et al. 2019) and particularly understudied when applied to non-regression frameworks such as the Markov models. In the context of discrete character models, the expected behavior is that rate estimates will tend towards zero, resulting in a downward bias of parameter estimates. Biologically, this means that a regularized model expects fewer changes in the character state through time and favors explanations of homology over homoplasy. The benefit of a regularized model is the increased generalizability should lead to a model which more accurately reflects the dynamics of unsampled taxa and leads to better predictions of unknown character states (e.g., ancestral character estimation).

To test the bias-variance trade-off associated with the regularization framework introduced here, I conduct a simulation study to evaluate the bias, variance, and overall error of parameter estimates for regularized and unregularized models. I simulate data under three model structures which vary only in the number of characters included. Specifically, I simulate data for a single character with binary states (k=2), two characters with binary states (k=8), and three characters with binary states (k=24). It was not required that all state combinations had to be observed. All models are simulated under an all rates different (ARD) model with dual transitions disallowed. Parameters are sampled from a log-normal distribution, where the logarithms of the parameters are normally distributed with a mean of 0 and standard deviation of 0.25. Parameters are sampled 100 times for a given phylogenetic tree. Phylogenetic trees of size 100, 250, and 500 taxa are simulated 10 times under a pure birth model with a sampling fraction of 1 using the R-package TreeSim (Stadler 2019). Phylogenies are then rescaled to a height of 1. These same 30 phylogenetic trees (10 each for the different number of taxa) are used for the 3 different model structures. In total, each model structure is tested under 3000 simulated datasets for a total of 9000 simulations.

Discrete character models are fit using the R-package corHMM (Beaulieu et al. 2022) with the simulating model structure (i.e., ARD) specified. Models are fit with no penalty term (unregularized), an *er* penalty term, an *l1* penalty term, and an *l2* penalty term with *λ*= 1. The maximum likelihood estimates are then compared to the simulating values and their bias, variance, and Root Mean Squared Error (RMSE) evaluated. Note that hidden Markov models are not included in Simulation Study I because, for trees of this size, it would be difficult to simulate data which consistently had a strong signal of rate heterogeneity (Boyko and Beaulieu 2021). This means that regularized models would show an inflated negative bias as parameters in a second-rate class would be estimated as 0 when there is weak evidence of hidden rates.

### Simulation Study II: Automatic model selection

Apart from reducing the variance of discrete character models, a primary aim of the dredge framework is to help users navigate large and complex model spaces. As such, in *Simulation Study II* I simulate data under three historically significant and widely applied model structures and evaluate whether the dredged model captures the most important features of the generating models. To conduct this evaluation, I compare three model selection strategies. The first is a standard unregularized greedy search, which acts as a baseline. The other two methods employ *1l* regularization to penalize model complexity, one using a greedy search algorithm and the other using a more exhaustive simulated annealing approach. Specifically, I simulate data under: (1) a correlated model, which captures the influence of two or more traits on each other (Pagel 1994); (2) an ordered and unidirectional model of trait change (e.g., Skinner et al. 2008); and (3) a hidden Markov model, where there is an unobserved factor causing rate variation in an observed character (Felsenstein and Churchill 1996; Beaulieu et al. 2013; Boyko and Beaulieu 2021). Each of these models has a unique set of criteria for the “most important features,” depending on what is typically considered relevant in empirical hypotheses using the particular model structure. Below I go into detail about which features are considered relevant for this study in each case.

When the generating model is consistent with Pagel’s (1994) correlated model, 100 datasets are simulated under *Q*_*corr*_ with transition rates of 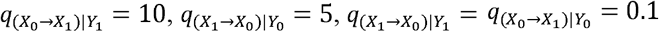 and 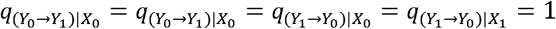. A unique pure birth phylogeny (birth rate = 1 events/MY) with 250 tips is simulated for each dataset and rescaled to have a height of 1. To evaluate whether the dynamics of the correlated model are adequately captured, I quantify how often does the best fitting dredge model find that: 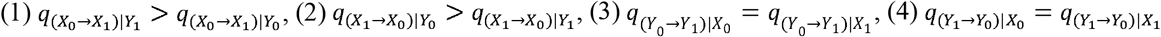, and (5) all other tests are passed. These comparisons are chosen because they are relevant to the interpretation of the correlated model. Evidence for the correlated model comes from tests of whether changes in the focal character depend on the state of the background character (Pagel and Meade 2006). As a hypothetical example, consider the situation in which *x* is parity mode and *Y* is climatic preference in reptiles: *x*_O_ is oviparity, *x*_1_ is viviparity, *Y*_O_ is a warm climatic preference, and *Y*_1_ is a cold climatic preference. An important factor for the evolutionary transition from egg-laying (oviparity) to live-birth (viviparity) in reptiles is the climate a lineage occupies, with squamate viviparity evolving frequently in colder climates, such as those found at high altitudes and latitudes (Blackbum 1999). When a correlated model of discrete character evolution is applied to this hypothetical example, the simulation parameters outlined above indicate that the transition of oviparity to viviparity is faster when a lineage is in a cold environment 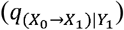 than when the lineage is in a warm environment 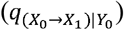 and that transitions from viviparity to oviparity are faster when lineages are in a warm environment 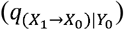 than when they are in a cold environment 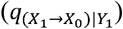. If parity mode evolved independently of climatic environment, then the expectations is that the transition rates between oviparity and viviparity are the same regardless of which environment the lineages inhabit. If corHMMDredge consistently finds 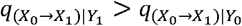 and 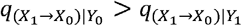, then it would seem to adequately capture correlated trait dynamics. Additionally, the simulation includes a negative control. For this simulation, transitions between climate do not dependent on the presence of viviparity or oviparity. This is captured by tests (3) and (4) examine whether transitions from warm to cold climates 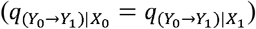 and cold to warm climates 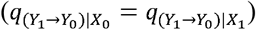 are the same regardless of the reproductive mode.

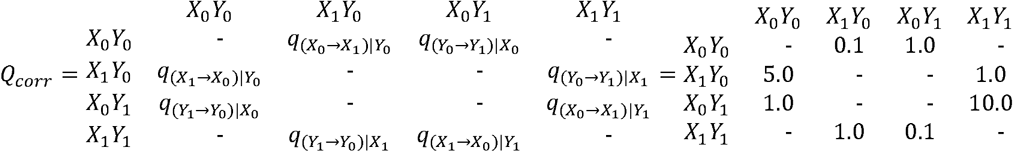

To simulate under an ordered trait evolution model, I construct a matrix Q_ord_ which allows for transitions from state 1 to state 2 to state 3, where reversions are possible only in state 2. All transition rates are set to 1, such that *q*_1 → 2_ = *q*_2 → 1_ = *q*_2 → 3_ =1 and a unique pure birth tree with 250 taxa is simulated with birth rate 1 for 100 datasets and trees are rescaled to a height of 1. To evaluate whether the dynamics of a ordered model are actually captured by the dredge framework, I conduct 4 tests: (1) *q*_1 → 3_ = 0, (2) *q*_2 → 1_ >0 ^ *q*_2 → 3_ >0, (3) *q*_3 → 1_ = *q*_3 → 2_ = 0, and (4) all other tests are passed. The first 3 tests examine whether the correct structure of the ordered model is discovered by corHMMDredge. A biological example of this type of model would be transitions between outcrossing and obligate selfing in angiosperms, through an intermediate facultative selfing state and the irreversibility of obligate selfing state. If this were the case, *x*_1_ would represent out-crossing, *x*_2_ would represent facultative selfing, and *x*_3_ would represent obligate selfing. The first test asks whether corHMMDredge has accurately discovered that it is impossible to directly transition from outcrossing to obligate selfing. The second test naturally follows the first examining whether transitions from the intermediate facultative selfing state are allowed to transition to both obligate selfing and outcrossing. The third tests asks whether the obligate selfing is correctly inferred to be a sink state, incapable of reverting back to either facultative selfing or outcrossing. This ordered model is somewhat unique among the historical models I am testing in that it is often clearly linked to apriori biological expectations related to developmental biology. And in those cases, it may not be appropriate to apply a dredge framework as the model set will be well defined. However, these sorts of ordered dynamics are possible in settings where developmental information is not readily available, and the goal of these simulations is to determine if the corHMMDredge framework can detect ordered models in general.

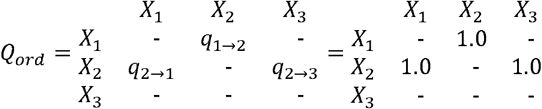

The final historically significant model being tested is a hidden Markov model where unobserved rate variation is introduced via a hidden factor. The matrix, *Q*_*HMM*_, describes a simple HMM for a single character, X, with two states, 0 and 1. The structure of an HMM is not unlike a correlated model, except that in the correlated case, both characters are observed and for an HMM we are trying to detect the hidden character based on the rate differences in the observed character (Boyko and Beaulieu 2021). For simulations, transition rates are set to be 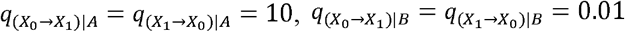, and 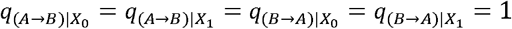 and simulated on 100 unique pure birth tree with 250 taxa. To evaluate whether the dynamics of a hidden Markov model are captured by the corHMMDredge framework, I conduct 2 tests: (1) 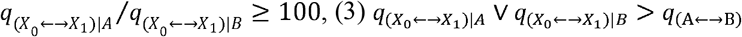.The first test asks whether there is support for hidden rates at all. The second test, conditioned on the first, asks whether the difference between the fastest rate class 100 times is at least greater than the slower rate class. I note that the simulating difference of 1000 times is a large difference (and arguably unrealistic) difference in rate classes but is chosen to ensure that simulating data had a signal of rate heterogeneity. Test 3 examines whether the fastest rates belong to transitions between observed states, rather than between hidden states.

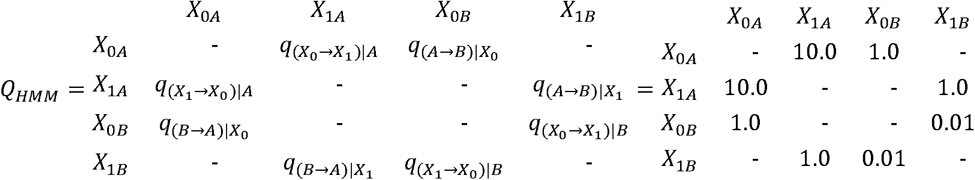

### Empirical example: concealed ovulation and mating systems in Old World monkeys

Concealed ovulation has been considered an adaptation in primates as either a means of promoting parental care by increasing confidence in paternity or improving male behavior towards their potential offspring by confusing paternity issues (Sillen-Tullberg and Moller 1993). Addressing whether concealed ovulation increases confidence or confusion in paternity issues is difficult to study at a phylogenetic scale, so previous examinations of this question have focused on the context in which the trait has evolved. An initial analysis by Sillen-Tullberg and Moller (1993) found that the common ancestor of all anthropoid primates showed signs of ovulation (or estrus advertisement) and had a multimale mating system. Concealed ovulation then evolved from this state 8 to 11 times in a nonmonogamous context and at most once in a monogamous context. In contrast, in their examination of this same question, Pagel and Meade (2006) found the common ancestor of all Old World primates to have concealed ovulation and a monogamous mating system. This discrepancy was explained by the difference in methodology. Pagel and Meade (2006) had used rate estimates from Markov models and estimating the ancestral state of the Old World primate common ancestor is crucial for the subsequent interpretation as it will imply whether estrus advertisement or concealed ovulation is the derived state in primates, though an updated dataset will likely be necessary before we can trust the robustness of any biological interpretations from these analyses.

To examine the evolution of concealed ovulation and mating systems, I apply corHMMDredge to the dataset of Pagel and Meade (2006). The dredge models use *l1* regularization and I conduct phylogenetic cross-validation to determine the optimal *λ*value. For cross-validation I examine 5 folds for *λ* values of 0, 0.25, 0.5, 0.75, and 1 with Jensen-Shannon divergence used as a metric of accuracy.

All model fits undergo 10 random restarts to increase the chances of finding the global optimum for the rate estimates. The best dredge model is used to estimate ancestral states based on the regularized maximum likelihood estimate. The root prior is set to “maddfitz” (FitzJohn et al. 2009) in the corHMM R-package. I examine the profile likelihoods of the rate estimates to generate 95% confidence intervals around the MLE independently for each parameter. To do this, I implement a new profile likelihood function within corHMM and examine 200 fixed values for each parameter spanning 100 orders of magnitude around the MLE. Profile likelihood operates by examining what the maximum likelihood would be given a fixed value and with the remaining parameters freely estimated.

## Results

### Simulation Study I: regularized models substantially reduce variance

The introduction of *l1* and *l2* regularization led to a significant reduction in total variance across all tested models for the raw parameter estimates (Table 1). Furthermore, the absolute average bias 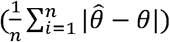 for phylogenies with 100 taxa and 250 taxa was found to be lower for *l1* and *l2* regularized models, contrary to the expected increase in negative bias (Table 1). This trend reversed for *l2* regularized models fit to datasets with trees of 500 taxa, but *l1* regularization consistently showed the lowest bias across all tested tree sizes. More surprisingly, the bias of *l1* regularized models was slightly positive on average, contrary to the expected downward bias, although a negative bias is found if one uses the median difference instead of the arithmetic mean. Finally, when considering overall error (RMSE), *l1* and *l2* regularized models showed substantial improvements over unregularized (*l0)* and *er* regularized models (Table 1). This improvement in RMSE highlights the effectiveness of *l1* and *l2* regularization in achieving better generalizability and prediction accuracy, despite the potential trade-off with increased bias. It is important to note that several of these findings go against expectation. Particularly, the absolute bias of an unregularized model is not expected to be greater than that of regularized models. There are two possible explanations for this which we can examine. First, it could be that several of the parameter estimates lack information to be well estimated causing them to reach an upper or lower bound during the maximum likelihood search. Indeed, as one removes outlier transition rate estimates the overall variance of the unregularized model estimates decreases substantially (Table 1). The second explanation is that parameter estimates should be evaluated on a log scale because fold changes are more meaningful for rate estimates than the raw units.

**Table 1.** Bias, variance, and overall error (RMSE) for regularized and unregularized models across phylogenies of varying sizes. Results are summarized across all generating models and presented in both original units and log-transformed units. Values in brackets are the bias, variance, or RMSE after filtering outliers. For original units, outliers are defined as having a transition rate estimate that is 10 transitions per unit time greater than the generating value. For log units, outliers are defined as being 5 orders of magnitude greater or lower than the generating value.

| Reg. type | Ntax a | Original units |  |  | Log units |  |  |
| --- | --- | --- | --- | --- | --- | --- | --- |
|  |  | Bias | Variance | RMSE | Bias | Variance | RMSE |
| *l0* | 100 | 1.65 (0.34) | 87.83 (3.92) | 9.45 (1.92) | -1.77 (0.09) | 15.29 (0.85) | 4.22 (0.68) |
| *l0* | 250 | 0.48 (0.23) | 15.91 (2.48) | 3.92 (1.43) | -1.08 (0.02) | 9.19 (0.57) | 3.11 (0.42) |
| *l0* | 500 | 0.16 (0.13) | 2.69 (1.52) | 1.45 (0.99) | -0.57 (0.002) | 4.95 (0.47) | 2.17 (0.29) |
| *er* | 100 | 1.68 (0.34) | 90.66 (3.94) | 9.6 (1.92) | -1.77 (0.09) | 15.32 (0.83) | 4.22 (0.67) |
| *er* | 250 | 0.47 (0.22) | 15.47 (2.5) | 3.87 (1.43) | -1.1 (0.02) | 9.42 (0.57) | 3.16 (0.42) |
| *er* | 500 | 0.16 (0.13) | 2.71 (1.52) | 1.46 (0.99) | -0.57 (0.003) | 4.86 (0.47) | 2.15 (0.29) |
| *l1* | 100 | -0.05 (-0.05) | 2.08 (2.03) | 1.4 (1.38) | -2.51 (-0.18) | 17.46 (0.65) | 4.82 (0.64) |
| *l1* | 250 | 0.04 (0.04) | 1.71 (1.66) | 1.18 (1.16) | -1.31 (-0.09) | 9.93 (0.51) | 3.32 (0.41) |
| *l1* | 500 | 0.04 (0.04) | 1.21 (1.18) | 0.86 (0.84) | -0.65 (-0.04) | 5.19 (0.43) | 2.25 (0.28) |
| *l2* | 100 | -0.5 (-0.5) | 0.46 (0.46) | 0.98 (0.98) | -2.81 (-0.38) | 16.43 (0.42) | 4.89 (0.7) |
| *l2* | 250 | -0.33 (-0.33) | 0.5 (0.5) | 0.81 (0.81) | -1.61 (-0.21) | 10.04 (0.34) | 3.48 (0.43) |
| *l2* | 500 | -0.21 (-0.21) | 0.49 (0.49) | 0.64 (0.64) | -0.86 (-0.12) | 5.57 (0.34) | 2.41 (0.32) |

When bias and variance are examined on log-transformed parameter estimates, I find that variance is slightly higher for regularized models than the unregularized models. Additionally, all models demonstrate a negative bias suggesting that transition rates are generally underestimated. Both bias and variance decrease as sample size increases. Once again, these results are counterintuitive, not only because the regularized models have a larger variance estimate, but because unregularized models are showing a consistent negative bias. Because I expect that this is due to outlier estimates I applied an aggressive filtering strategy, removing parameter estimates that 5 orders of magnitude above or below the true generating parameter values (Table 1; for original units the filtering strategy is to remove differences in transition rates of greater than 10 units). Unsurprisingly, without outliers the bias, variance, and RMSE all decrease substantially. More importantly, when examining log-transformed parameter estimates, I find that regularized models have a negative bias with an absolute bias that is greater than the unregularized model and that regularized models have a lower variance than the regularized model matching expectations(Table 1).

### Simulation Study II: performance for model selection

Both regularized and unregularized methods preformed equally well when it came to automatically inferring the generating model structure (Table 2). The most substantial differences are seen when comparing the simulated annealing routine versus a greedy hill climbing approach (9 out of the 12 tests simulated annealing preforms as well or better). For the correlated model, while all methods accurately recovered the primary dependencies, the simulated annealing search was roughly twice as successful at passing all focal tests simultaneously (10.1% vs 5.1-6.1%). Its advantage was more pronounced for the ordered model, where simulated annealing was substantially more effective at identifying both forbidden transitions and irreversible sink states, leading to an overall success rate of 33% compared to ~20% for the greedy searches. Finally, for hidden states model, simulated annealing was better at detecting the large magnitude difference between the fast and slow rate classes, succeeding in 75.8% of simulations compared to 65.3% for the *l1-*regularized greedy search and only 25.3% for the unregularized greedy search. This indicates that while greedy searches are generally proficient, the more exhaustive search of simulated annealing can help uncover additional complex model structures, particularly those involving rate heterogeneity.

**Table 2.** Results from structure tests comparing a greedy hill climbing search to simulated annealing (SA). Right hand values indicate the proportion of time each framework passed the row’s Focal Test. All methods performed reasonably in identifying the generating model structure.

| Model | Focal Test | Greedy (l0) | Greedy (l1) | SA (l1) |
| --- | --- | --- | --- | --- |
| dependent | $q_{(X_0 \rightarrow X_1) \vert Y_1} > q_{(X_0 \rightarrow X_1) \vert Y_0}$ | 0.98 | **0.99** | 0.97 |
| dependent | $q_{(X_1 \rightarrow X_0) \vert Y_0} > q_{(X_1 \rightarrow X_0) \vert Y_1}$ | 0.899 | 0.899 | **0.909** |
| dependent | $q_{(Y_0 \rightarrow Y_1) \vert X_0} = q_{(Y_0 \rightarrow Y_1) \vert X_1}$ | **0.303** | 0.293 | **0.303** |
| dependent | $q_{(Y_1 \rightarrow Y_0) \vert X_0} = q_{(Y_1 \rightarrow Y_0) \vert X_1}$ | 0.162 | 0.253 | **0.293** |
| dependent | All dependent model tests passed | 0.051 | 0.061 | **0.101** |
| ordered | $q_{1 \rightarrow 3} = 0$ | 0.66 | 0.54 | **0.71** |
| ordered | $q_{2 \rightarrow 1} > 0 \wedge q_{2 \rightarrow 3} > 0$ | **0.53** | 0.48 | 0.51 |
| ordered | $q_{3 \rightarrow 1} = q_{3 \rightarrow 2} = 0$ | 0.44 | 0.68 | **0.72** |
| ordered | All ordered model tests passed | 0.21 | 0.2 | **0.33** |
| hidden states | $n\_rate\_class > 1$ | 0.726 | 0.863 | **0.905** |
| hidden states | $q_{(X_0 \leftrightarrow X_1) \vert A} / q_{(X_0 \leftrightarrow X_1) \vert B} \geq 100$ | 0.253 | 0.653 | **0.758** |
| hidden states | $q_{(X_0 \leftrightarrow X_1) \vert A} \vee q_{(X_0 \leftrightarrow X_1) \vert B} > q_{(A \leftrightarrow B)}$ | 0.705 | **0.8** | 0.789 |

### Canalization of mating systems through time

The ancestral state reconstruction at the root under the dredge model had considerable uncertainty regarding the evolutionary origins of mating system traits. The most likely estimated ancestral state was concealed ovulation with a monogamous mating system (state 00) having a marginal probability of 61%, while state 01 (estrus display and monogamous mating system) had a marginal probability of 38% (Fig. 3a). The result of simulated annealing identified a two-parameter model incorporating hidden rates as the optimal framework for describing trait evolution (Fig. 3b). The parameter estimates revealed distinct transition rate classes: states 00, 01, and 11 shared a common transition rate of 0.038 transitions/MY (95% CI: 0.017, 0.093), while transitions from 00R1 to 00R2 and 11R1 to 11R2 occurred at a rate of 0.029 (95% CI: 0.00099, 0.12). A particularly notable feature of the optimal model is the presence of two sink states within the R2 rate classes. Once a lineage transitions to rate class two for either concealed ovulation with a monogamous mating system with (00R2) or estrus display with multimale mating (11R2), the model predicts no further evolutionary transitions, effectively trapping these lineages in their current state. This is similar to the covarion model, where molecular evolutionary processes can cease entirely in certain lineages (Tuffley and Steel 1998). However, this model differs in that lineages entering these sink states are not expected to resume evolutionary transitions, suggesting a form of evolutionary canalization rather than temporary stasis. Based on state reconstructions at the tips for the unknown hidden states, there are some clade-specific patterns of evolutionary canalization. Clade 1 (*Hylobates*) appears to have entered this canalized state, remaining permanently as monogamous mating with concealed ovulation (00), while Clade 2 (*Macaca*) is trending towards a canalization of multimale mating and estrus display (11). Additionally, the common ancestor of humans and their closest relatives is estimated to be at an intermediate state (01), which subsequently diverged to state 00R1 in humans and state 11R1 in both *Pan paniscus* and *Pan troglodytes* (Clade 3).

**Figure 3.**
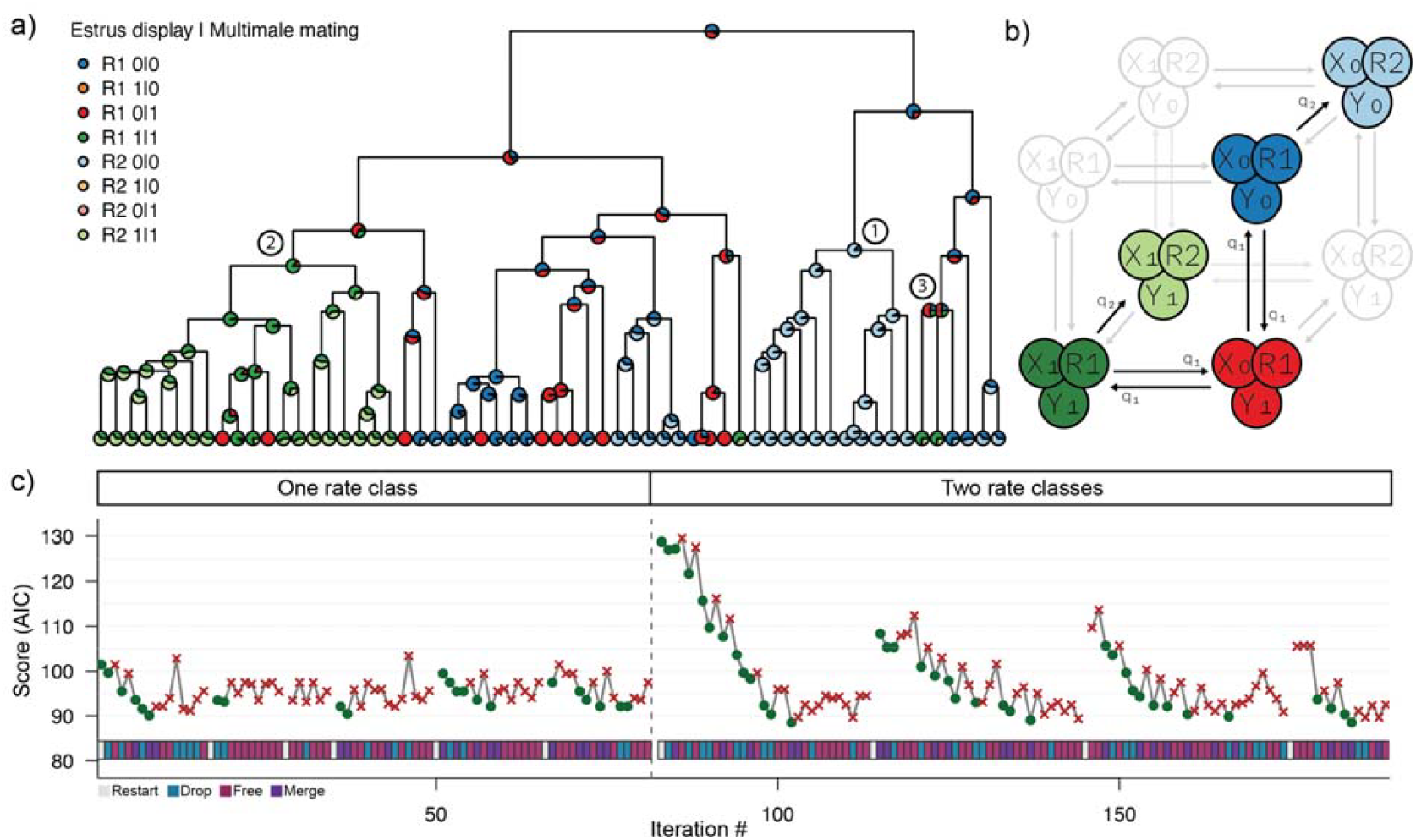
Results of the empirical dredge analysis of primate mating systems and ovulation. (a) Ancestral state reconstruction on the phylogeny under the best-fitting model. Node and tip colors correspond to the combined character states and hidden rate classes (R1, R2). Numbered clades are referenced in the text. (b) The structure of the final dredged model, a two-parameter hidden Markov model. Colored states and solid arrows represent the inferred model, while grayed-out elements show the full potential model space. Note the “sink” states (00R2 and 11R2) which have no outgoing transitions. (c) The simulated annealing trace plot, showing the AICc score over iterations for models with one and two rate classes. The bar at the bottom indicates the type of move proposed at each step.

The dredge analysis was conducted using a temperature of 4 and cooling rate of 0.9 (Fig. 3c). However, the initial run’s final model included a transition involving an unobserved state (10), suggesting that the simulated annealing routine had not reached the global optimum and the temperature parameter maybe have been too low for the model’s complexity. To address this limitation, a second dredge run was initiated using the first run’s final model as a starting point. This iterative approach demonstrates the utility of human-guided model refinement in corHMMDredge, allowing researchers to apply biological intuition to improve model realism. Finally, cross-validation analysis supported the robustness of the final model, with a lambda value of 1 yielding the highest validation scores (0.2234; Table 3). While our cross-validation was performed on the final dredged model, I note that this approach may introduce a bias, as the model search itself was conducted under a fixed lambda. Ideally, cross-validation would be integrated into the entire simulated annealing routine. However, given the significant computational cost of this procedure, I recommend most users perform cross-validation as a final, confirmatory step. For studies where computational resources are not a limiting factor, running the entire dredge search across a range of lambda values and comparing the cross-validation scores of each final model would provide the most rigorous validation.

**Table 3.** Results of the 5-fold cross-validation used to determine the optimal l1 regularization penalty (lambda) for the final dredged model. Each value represents the validation score for a given fold and lambda setting. The optimal value was determined to be lambda = 1.0, as it produced the best average score across all five folds.

| **Lambda** | **0** | **0.25** | **0.5** | **0.75** | **1** |
| --- | --- | --- | --- | --- | --- |
| *Fold 1* | 0.2836 | 0.2259 | 0.2256 | 0.2254 | 0.2252 |
| *Fold 2* | 0.2722 | 0.2180 | 0.2171 | 0.2163 | 0.2156 |
| *Fold 3* | 0.2282 | 0.2276 | 0.2271 | 0.2266 | 0.2262 |
| *Fold 4* | 0.2626 | 0.2617 | 0.2608 | 0.2600 | 0.2592 |
| *Fold 5* | 0.1918 | 0.1915 | 0.1913 | 0.1912 | 0.1911 |
| *Average Score* | 0.2477 | 0.2249 | 0.2244 | 0.2239 | 0.2235 |

## Discussion

### Uncertainty in ancestral estimation and the challenges of model interpretation

The dredge algorithm was able to find a unique, non-standard, model structure in the empirical case study (Fig. 3b). The dredge model estimated the ancestral state at the root to be a monogamous mating system with concealed ovulation (marginal probability of 61%), though it was fairly uncertain. Sillen-Tullberg and Moller (1991) found that the root state was likely to be estrus advertisement with multimale mating system and therefore discussed the evolution of concealed ovulation in this context. In contrast, and as in this study, Pagel and Meade (2006) had found that the best supported root state was concealed ovulation with monogamous mating. In the absence of direct observation, it is nearly impossible to say which of these outcomes is more correct. However, revisiting the hypotheses of Sillen-Tullberg and Moller (1991) under the dredged model results do provide some interesting alternative interpretations for the evolution of estrus advertisement and mating system.

Sillen-Tullberg and Moller (1991) had considered concealed ovulation to be the derived state and found that it had almost always evolved in non-monogamous lineages. This provided evidence that concealed ovulation was more likely to be a prerequisite for the evolution of monogamy, rather than a consequence of it (as had been hypothesized by (Burley 1979). However, these modeling results suggest an entirely different pathway with estrus display evolving only after multimale mating has established itself. This finding somewhat contradicts the hypothesis that signs of ovulation will disappear in non-monogamous mating systems because paternity confusion will be beneficial (Hrdy 1979). That is one possible outcome and is inferred to occur at a rate of 0.038 transitions/MY (11 → 01), but from the concealed ovulation and multimale state, a lineage is just as likely to regain overt ovulation signs as it was to have lost them in the first place (0.038 transitions/MY; 01 → 11). This is because under the dredged model structure, concealed ovulation with multimale mating, is an intermediate state. Furthermore, a particularly novel insight from the dredged model is the inference of evolutionary canalization. The model identifies sink states where, after a lineage transitions into a second rate-class for either monogamy and concealed ovulation (00R2) or multimale mating with estrus display (11R2), it becomes evolutionarily trapped. Though it is not possible to know the mechanism from comparative data, this model could explain the relative stability of mating systems in certain clades like gibbons.

There are two important caveats when interpreting these results. First, this biological interpretation is contingent on the single best-fitting model. While this model had the lowest AIC score, several others had values well within the recommended threshold of ΔAIC < 2 (Burnham and Anderson 2002), implying that alternative evolutionary scenarios also have substantial support. As discussed in the section Choosing the “best” model, model averaging to account for this uncertainty is non-trivial when dealing with hidden states. Second, any comparative analysis is fundamentally dependent on the input data. This study re-analyzes the Pagel and Meade (2006) dataset, which is now nearly two decades old. Revisions to the phylogeny or character state assignments in that time could significantly alter the outcome. Therefore, while the dredged model provides a compelling alternative hypothesis for the evolution of mating systems and ovulation, its conclusions should be viewed as provisional and should be reexamined with contemporary data.

### The cost of regularization and eliminating very high transition rates

The dredge algorithm, and regularization more generally, will result in a downward bias proportional to the magnitude of the transition rate. It is worth considering what, if any, is the cost of this downward bias. In empirical settings, it is not uncommon to have transition rates approaching and being estimated at an upper bound. When interpreted as 100s of state transitions occurring over short time intervals, this may be cause for concern because it often seems biologically implausible. However, high transition rates do not necessarily predict that many transitions of that type have occurred. Often, high transition rates may correspond to only a few inferred transitions on a tree (though they will have occurred very rapidly). The two inferences, extremely high transition rates and relatively few transitions, may seem to be contradictory, but they are perfectly compatible since a lineage must first be in the initial state to transition, and the initial state need not be common. Instead, a more serious cause for concern is the amount of information available to infer the transition rate if that transition only occurs rarely throughout the clade’s history. This was the case for the Old World monkey dataset when examining an all-rates-different model. Several parameter estimates seemed to have very little information and fell along a likelihood ridge. The combination of regularization and model structure searching removes this ridge but also has the benefit of estimating rates which better match intuition that a transition which occurs rarely throughout the clade has a corresponding transition rate that is also low. Ultimately, a regularization scheme moves towards a more parsimonious view of character evolution, with rate estimates tending to be biased towards explanations of homology over homoplasy.

### The value of multiple characters

Part of the value of a dredge approach lies in its ability to identify important relationships between characters by eliminating or equating parameters that lack significant support. This is particularly useful for phenotypic complexes with multiple interacting characters (e.g., pollinator syndromes). However, a dataset containing many characters also introduces a large state space with numerous testable model structures. One solution to address this issue is to independently model each character and then compile the results into a cohesive view. This approach is computationally tractable and can yield well-behaved parameter estimates. But modeling each character independently ignores correlated character evolution and dependent relationships can influence both biological inferences and ancestral state estimation (Boyko and Beaulieu 2021). When the evolution of one character affects the rate of change in another, independent modeling may lead to uncertainty in certain parts of the phylogeny. By accounting for correlated character evolution, shared information between characters can improve inference (Boyko and Beaulieu 2021). Of course, correlation between discrete characters is a hypothesis that should first be tested. These tests should account for character-independent rate variation (Boyko and Beaulieu 2023), which will introduce even more parameters than standard independent models (Pagel 1994). Searching a state space that includes multiple characters and hidden rate classes can be a daunting task even for experienced comparative biologists and it can be tempting to rely on the most used default model sets. However, there is greater flexibility in potential dependent relationships beyond typical fixed model sets. Correlation tests have traditionally compared independent and correlated models, but not all characters need to show dependent relationships for there to be evidence of correlation (see also Pagel and Meade 2006). A dredge framework is valuable here, as it searches potential model structures and identifies only the necessary dependent relationships.

### Choosing the “best” model

Model averaging is central to both Bayesian frameworks (e.g., Pagel and Meade 2006) and frequentist multi-model inference (MMI; Burnham and Anderson 2002; Jhwueng et al. 2014) and improves predictive power by integrating results over a set of plausible models. In contrast, model selection aims to identify a single best-fitting model or a small set of top models, often guided by an information criterion that balances model fit and complexity. While model averaging is powerful, its direct application following the use corHMMDredge faces some technical challenges, leading to a methodology centered on model selection and examination of the top-ranked models. The most critical challenge is the label switching problem inherent to hidden Markov models (Stephens 2000). For example, one model might assign high transition rates to hidden rate class 1, while another equally likely model assigns them to rate class 2. Averaging parameters across these models without resolving this arbitrary class assignment would produce meaningless estimates and incorrect conclusions. This issue can be mitigated in some cases, such as in diversification models, by focusing on tip rates rather than internal parameters (Title and Rabosky 2019; Vasconcelos et al. 2022).

However, a further complication with model averaging is the potential for pseudo-replication of model structures. The model space for more complex models (e.g., those with two rate classes or correlated character evolution) is inherently larger than that for simpler models (e.g., with one rate class or independent evolution). As a result, the total number of models available for averaging may disproportionately represent more complex models, even when there is roughly equal support for simpler alternatives. This can distort model averaging, as the AIC weights may reflect the sheer number of models rather than the actual support for distinct, biologically meaningful models. The most extreme example of this would be if identical models are refit, effectively double counting them. However, this problem can also occur if models are nearly identical, such as when two parameters are equated (e.g., k=7 vs. k=8). These models might still receive significant support despite being only minor variations of the most complex model. Handling this model structure pseudo-replication becomes particularly important for automated model selection.

This was not a problem when Burnham and Anderson (2002) originally proposed multi-model inference because they emphasized fitting a set of biologically plausible (and distinct) models (Burnham and Anderson 2002). Each model was justified based on specific hypotheses and therefore tested the contribution of different factors. For exploratory analyses on large model spaces, the rationale behind model averaging becomes less clear. In a dredge run, one is not testing specific hypotheses but rather comparing parameter estimates across many models. The final model of a dredge run does not necessarily reflect the combination of several well-supported biological models but is the byproduct of optimizing the bias-variance trade-off through AIC and encouraging sparse parametrizations through regularization. This is valuable from an inference perspective since the more complete a model set is, the less likely it is for unanticipated biases to creep into the results (Maddison and FitzJohn 2015; Rabosky and Goldberg 2015).

Nonetheless, the dredge approach and MMI need not be mutually exclusive. In cases where a set of biologically plausible hypotheses are known beforehand, a MMI approach remains an excellent option. And so long as procedures are clearly documented, there are advantages to combining the approaches by considering which model structures are biologically reasonable and determining what the optimal model structure could be for a given dataset. Furthermore, the most robust biological inferences can then be drawn not by relying on a single “best” model, but by examining the top models in the corHMMDredge set (e.g., those within ΔAIC < 2). This will allow researchers to assess whether key parameter estimates and biological interpretations are stable across a range of well-supported model structures. And because these models are ultimately phenomenological, the biological insights we draw from the best-supported model, and how these insights shape our understanding of evolutionary processes, are ultimately derived from subsequent interpretations and considerations in concert with other pieces of evidence (Gardner and Organ 2021). This perspective leaves room for quantifying the patterns in a general way via comparative modeling results, without necessarily forcing too much biological meaning on any particular parameter estimate.

## Conclusion

Determining which models to test in an empirical setting is an incredibly valuable process in which biologists take hypotheses and formally structure them so that they can be compared. This process can help clarify thinking around biologically relevant interactions between variables and will lead to more robust inferences. However, if only a subset of model structures are considered (e.g., only examining all-rates-different and equal-rates), there is a risk of overlooking plausible and important model structures. This issue is exacerbated by the fact that the knowledge necessary to manipulate PCMs is often difficult to acquire and may be hidden in highly technical texts, making it challenging for biologists to explore and customize model sets effectively (Cooper et al. 2016). It is evident that a comprehensive model set is important for trustworthy inferences in comparative biology (FitzJohn et al. 2009; Rabosky and Goldberg 2015; Beaulieu and O’Meara 2016; Boyko and Beaulieu 2023), but the growing complexity of discrete character models makes it challenging for users to determine which models are potentially realistic and important to consider. Here I have shown how the corHMMDredge framework may help alleviate this burden. This framework will enable biologists to focus more on model interpretations rather than model construction and may even lead to the discovery of model structures which imply unique hypotheses that would have not been considered otherwise.

## Acknowledgements

The author thanks Thais Vasconcelos and the Smith lab for their feedback on the overall clarity of the manuscript. The author would also like to thank Brian O’Meara for his thoughtful comments on an earlier version of this manuscript and pointing out the similarity of this approach and Sanderson’s (2002) penalized likelihood. The author thanks Rudolf Schill and Yutong Wang for general discussions about regularization approaches. Finally, the author would like to thank Sebastian Höhna, Josef Uyeda, and two anonymous reviews for their thoughtful feedback which greatly improved the quality of this manuscript.

## Data Availability

All the original data and scripts necessary to reproduce the analyses reported in this study can be accessed through the Dryad link: http://datadryad.org/share/HmmoM5OiQz2KUQr230La7ntdprGKF3kUR5rU0N-CZjs

## Supplementary Material

Supplementary files including pseudocode for simulated annealing search and the relative model support for the Old World monkey analysis is available at SYSBIO online.

## Pseudocode for simulated annealing search

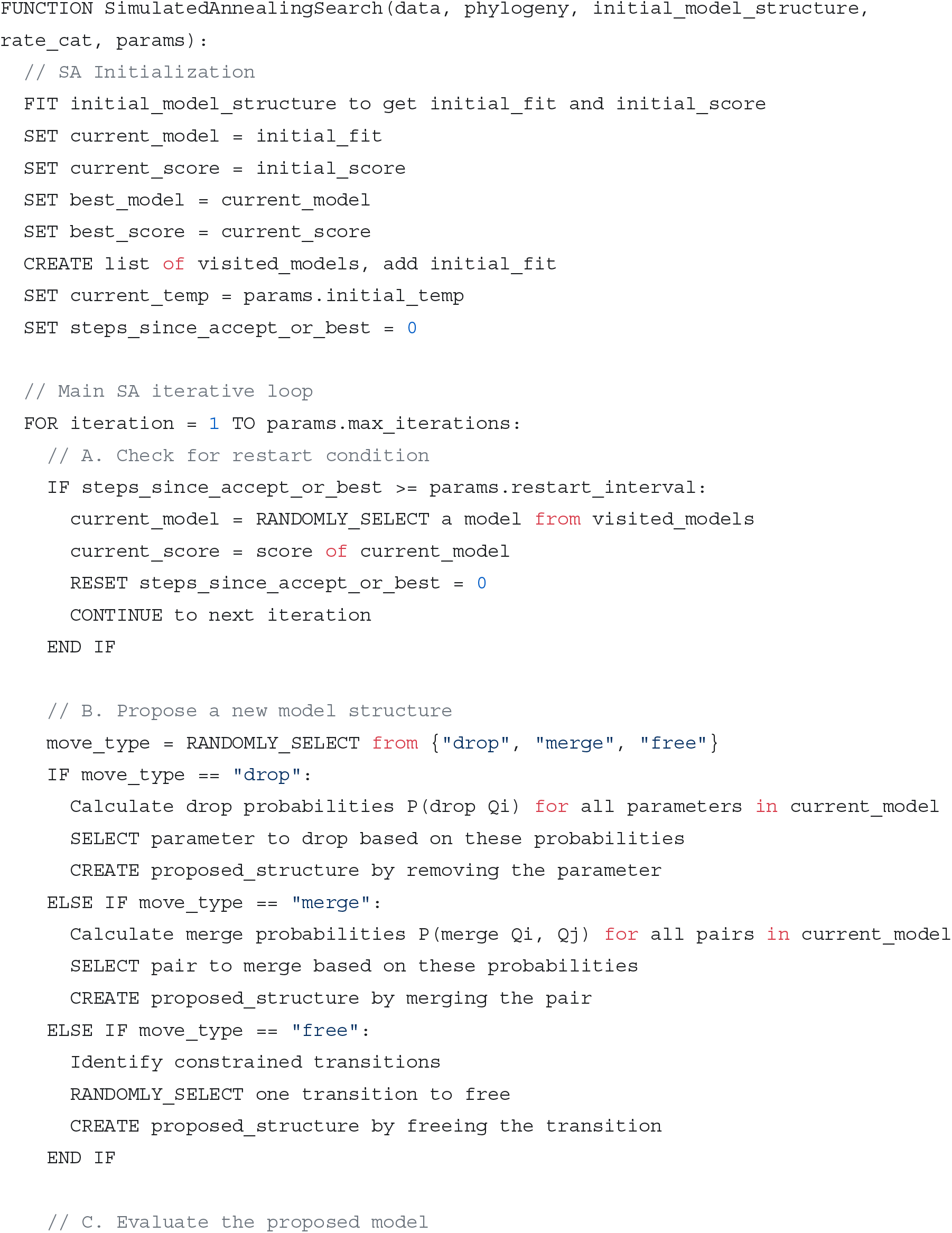
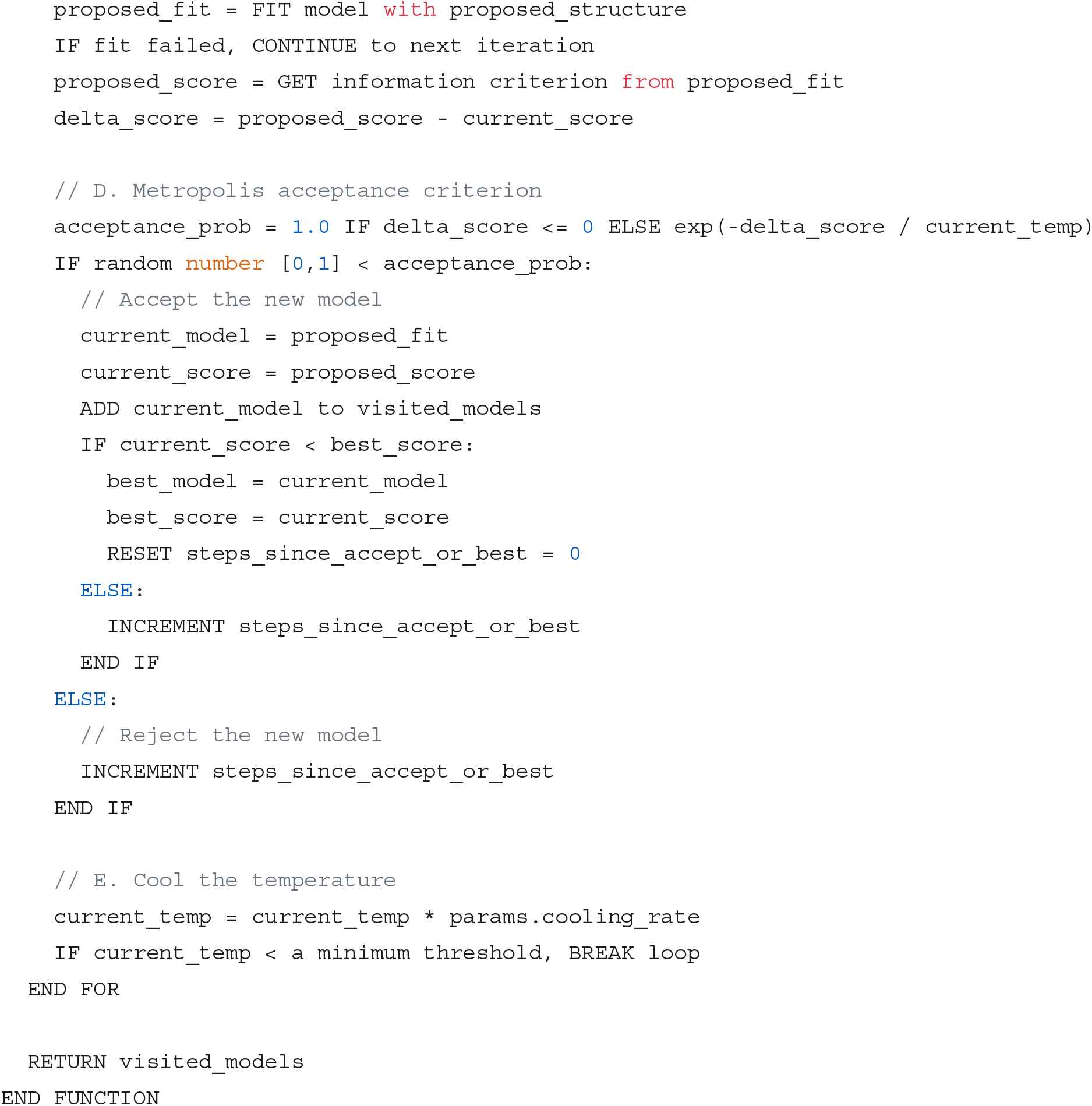

---

**Table S1.** Relative model support for the Old World monkey analysis. This table includes the number of parameters (np), the number of rate categories (nRateCat), the log likelihood value for a given model (lnLik), the Akaike Information Criterion value (AIC), the difference from the models AIC value to the best AIC value (dAIC) and the AIC weight. These are the unique models which were accepted during the simulated annealing routine. Many more models were attempted but not accepted during the search.

| np | nRateCat | lnLik | AIC | dAIC | AICwt |
| --- | --- | --- | --- | --- | --- |
| 4 | 1 | -42.756941 | 93.5138814 | 5.59242874 | 0.0084073 |
| 7 | 1 | -42.823622 | 99.6472448 | 11.7257921 | 0.00039157 |
| 3 | 1 | -42.791402 | 91.5828036 | 3.66135092 | 0.02207928 |
| 2 | 1 | -43.055279 | 90.1105582 | 2.18910557 | 0.04609766 |
| 3 | 1 | -43.570587 | 93.1411748 | 5.21972214 | 0.01012952 |
| 2 | 1 | -43.231785 | 90.4635702 | 2.54211759 | 0.03863877 |
| 24 | 2 | -40.360785 | 128.721571 | 40.800118 | 1.90E-10 |
| 23 | 2 | -40.505853 | 127.011707 | 39.090254 | 4.47E-10 |
| 22 | 2 | -41.620411 | 127.240823 | 39.31937 | 3.99E-10 |
| 6 | 2 | -40.839915 | 93.6798291 | 5.75837645 | 0.00773787 |
| 17 | 2 | -40.840757 | 115.681515 | 27.7600622 | 1.29E-07 |
| 14 | 2 | -40.83998 | 109.67996 | 21.758507 | 2.60E-06 |
| 4 | 2 | -41.181067 | 90.3621331 | 2.4406804 | 0.04064902 |
| 3 | 2 | -41.2575 | 88.514999 | 0.59354638 | 0.10236468 |
| 12 | 2 | -40.673954 | 105.347907 | 17.4264544 | 2.26E-05 |
| 12 | 2 | -40.673961 | 105.347921 | 17.4264684 | 2.26E-05 |
| 6 | 2 | -40.504566 | 93.0091319 | 5.08767929 | 0.01082085 |
| 8 | 2 | -40.948669 | 97.8973384 | 9.97588573 | 0.00093929 |
| 6 | 2 | -40.948663 | 93.8973261 | 5.97587344 | 0.00694053 |
| 5 | 2 | -41.189251 | 92.3785021 | 4.45704939 | 0.01483205 |
| 3 | 2 | -41.533733 | 89.0674668 | 1.1460141 | 0.07765748 |
| 11 | 2 | -40.839923 | 103.679846 | 15.7583936 | 5.21E-05 |
| 9 | 2 | -40.839917 | 99.6798338 | 11.7583811 | 0.00038525 |
| 7 | 2 | -40.839914 | 95.679829 | 7.75837634 | 0.0028466 |
| 4 | 2 | -42.052089 | 92.1041774 | 4.18272478 | 0.01701257 |
| 3 | 2 | -42.192072 | 90.3841433 | 2.46269067 | 0.04020412 |
| 3 | 2 | -41.935946 | 89.8718921 | 1.9504394 | 0.05194031 |
| 5 | 2 | -40.839919 | 91.6798375 | 3.7583848 | 0.02103363 |
| 3 | 2 | -41.175615 | 88.3512292 | 0.42977656 | 0.11109955 |
| 3 | 2 | -42.56508 | 91.130161 | 3.20870832 | 0.02768694 |
| 2 | 2 | -43.114338 | 90.2286759 | 2.30722325 | 0.04345402 |
| 3 | 2 | -41.300881 | 88.6017622 | 0.68030953 | 0.09801888 |
| 3 | 2 | -42.779014 | 91.5580279 | 3.63657528 | 0.02235449 |
| 3 | 2 | -43.381054 | 92.7621085 | 4.8406558 | 0.0122434 |
| 4 | 2 | -41.620225 | 91.240449 | 3.31899638 | 0.0262015 |
| **2** | **2** | **-41.960726** | **87.9214527** | **0** | **0.13773271** |

